# Presynaptic occlusion of NMDA receptor-dependent potentiation in Synaptophysin family knockouts

**DOI:** 10.1101/2024.03.12.583943

**Authors:** Sergio del Olmo-Cabrera, Juan José Rodríguez Gotor, Rémy Verhaeghe, John F. Wesseling

**Affiliations:** Institute for Neuroscience, CSIC-UMH, San Juan de Alicante, Spain

**Keywords:** LTP, Long-term potentiation, synaptophysin, Syp, synaptogyrin, Syngr1, Syngr3

## Abstract

Baseline synaptic strength is elevated in knockouts missing the synaptophysin family of synaptic vesicle proteins because of increased probability of release of release-ready vesicles. Here we show that the longest lasting component of long-term potentiation (LTP) is: (a) eliminated at Schaffer collateral synapses under standard conditions; (b) rescued by reducing extracellular Ca^2+^ enough to lower baseline synaptic strength to wildtype levels; and, (c) eliminated at wildtype synapses by increasing Ca^2+^ enough to elevate baseline to knockout levels. Next, an earlier component of LTP, lasting less than 1 hr, was eliminated first at knockouts and then wildtype by elevating baseline further. And, the LTP that was eliminated could be recovered by subsequently reducing extracellular Ca^2+^ back to the permissive level. In sum, NMDA receptor-dependent LTP maintenance and expression are separable mechanisms at extensively studied Schaffer collateral synapses, and expression of multiple components can be occluded by enhancing the mechanism that catalyzes synaptic vesicle exocytosis from presynaptic terminals.

## Introduction

Synaptophysin family proteins are ubiquitously expressed within synaptic vesicle membranes at high copy number; family members in mammalian synapses are Synaptophysin (Syp), Synaptoporin (Synpr), and Synaptogyrins 1 (Syngr1) and 3 (Syngr3) [1, 2]. Despite the abundance, family members are not required for high fidelity synaptic transmission. Indeed, to the contrary, the *probability of release* of *readily releasable vesicles* – denoted with *p_v_* – is elevated in knockout synapses missing all four neuronal family members (SypFamilyKO) [3].

Readily releasable vesicles are thought to be docked to *release sites* embedded within the active zone of the plasma membrane of presynaptic terminals, and the elevated *p_v_* implies that sites at SypFamilyKO synapses catalyze exocytosis more efficiently than wildtype sites. In contrast, no alterations were detected at SypFamilyKO synapses in other basic parameters of synaptic vesicle trafficking such as the number of readily releasable vesicles or the timing of rate-limiting steps during sustained high frequency stimulation [3]. These results and others where exocytosis in PC12 cells was inhibited by heterologous expression of Syp and Syngr1 suggest that endogenous synaptophysin family proteins are negative regulators of the mechanism that catalyzes exocytosis [4]. However, deficits have additionally been observed in both short– and long-term plasticity of synaptic strength at Schaffer collateral synapses from knockout mice missing only two of the four family members [5], and it was unclear if the deficits in plasticity are related to the elevated *p_v_*.

Both short– and long-term plasticity have been subdivided into components that can be distinguished by the timing of reversion to baseline after activation. Here we investigated the relationship between the elevated *p_v_* at SypFamilyKO synapses and both the quickly reverting component of short-term plasticity termed *facilitation* – sustained for less than 1*/*2 s – and slowly reverting components of long-term plasticity (LTP) that are sustained for tens of minutes to hours, which is *>* 4 orders of magnitude (10 000-fold) longer.

A key for understanding our results is the distinction between *induction*, *maintenance*, and *expression* [6]. Induction is the event that activates the plasticity. Maintenance is the mechanism that sustains the plasticity, and determines the timing of reversion of each component to baseline. And, expression is the mechanism that physically realizes the change in synaptic strength.

The previous evidence for elevated *p_v_* at SypFamilyKO synapses pertains primarily to baseline whereas activity dependent modulation of *p_v_* was not investigated thoroughly. Even so, we hypothesized previously that the elevated baseline *p_v_* at SypFamilyKO synapses causes a deficit in facilitation by occluding expression [3]. That is, facilitation is: induced by the Ca^2+^ remaining within presynaptic terminals after action potentials; maintained until the Ca^2+^ is cleared; and expressed via increased *p_v_* (e.g., [7]). And, a variety of factors that increase *p_v_* are already known to occlude expression of facilitation at wildtype synapses because of a still poorly understood ceiling on the maximum value for *p_v_* [8]. Because of this, we reasoned that baseline *p_v_* might be closer to the ceiling in SypFamilyKO synapses, which would then occlude facilitation by restricting the remaining dynamic range. And indeed, supporting the hypothesis: facilitation was less at SypFamilyKO synapses compared to wildtype under standard conditions, but could be increased by lowering extracellular Ca^2+^ [3], which would relieve any occlusion by decreasing baseline *p_v_* owing to less Ca^2+^ flux into presynaptic terminals during action potentials [9–13].

LTP at Schaffer collateral synapses is more complicated than facilitation because induction requires: glutamate neurotransmitter release from presynaptic terminals; activation of NMDA-type glutamate receptors embedded within the membrane of postsynaptic compartments of separate neurons; and then second messenger signaling within the postsynaptic compartments [14]. However, previous studies have suggested that at least one component of LTP might be expressed by presynaptic mechanisms activated by retrograde signals emitted by the postsynaptic compartment [15–21]. And indeed, elevated baseline *p_v_* at neonatal synapses likely occludes LTP [22]. Here we report that, like facilitation, the longest-lasting component of LTP expression at mature SypFamilyKO synapses is occluded by the elevated baseline *p_v_*, and that an early component can be occluded by elevating the baseline even more; mechanisms of induction and maintenance were not affected.

## Results

### Expression of facilitation is occluded at SypFamilyKO synapses

The hypothesis that expression of facilitation is occluded by the elevated baseline *p_v_* at SypFamilyKO synapses predicts that the amount expressed at SypFamilyKO and wildtype synapses would be equal when baseline *p_v_* is lowered enough to relieve the occlusion completely. The prediction was not tested in the previous study where lowering extracellular Ca^2+^ increased paired-pulse facilitation at SypFamilyKO synapses [3], leaving open the alternative possibility that the deficit at SypFamilyKO synapses is caused by a partial defect in induction rather than by occlusion of expression. We now confirm: that paired-pulse facilitation is less at SypFamilyKO synapses under standard conditions where Ca^2+^ is 2.6 mM (Fig 1a); that decreasing Ca^2+^ to 0.5 mM greatly decreases synaptic strength (Fig 1b); and that facilitation at SypFamilyKO synapses equals the amount at wildtype when Ca^2+^ is 0.5 mM (Fig 1c-e).

**Figure 1.**
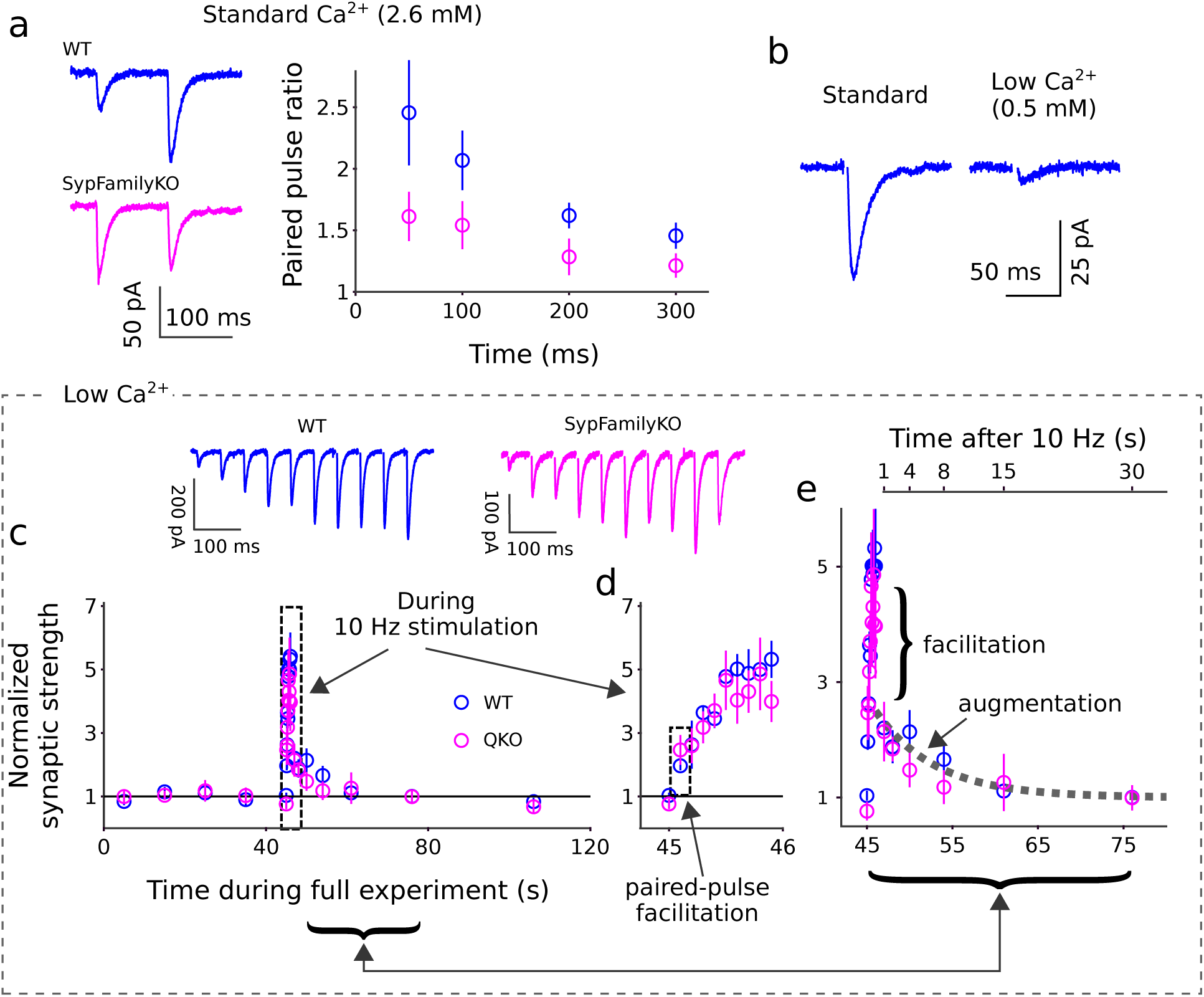
No deficit in short-term enhancement in low Ca^2+^. **(a)** Less paired-pulse facilitation at SypFamilyKO synapses under standard conditions. Traces are patch clamp recordings from individual trials after blanking the stimulus artifact. Paired pulse ratio is the current integral of the second divided by first response (*n* 5 neurons; experiments for each time point were repeated 6 times for each neuron and then digitally averaged before measuring). Ca^2+^ and Mg^2+^ were 2.6 and 1.3 mM. **(b)** Example of large decrease in response size when reducing Ca^2+^ to 0.5 mM (Mg^2+^ was 1.5 mM). **(c-e)** Experiments in 0.5 mM Ca^2+^. Schaffer collaterals were stimulated 5 times, once every 10 s, to acquire a baseline, then with 10 pulses at 10 Hz to induce short-term enhancement, and finally with single pulses 1, 2, 4, 8, 15, 30, and 60 s after the end of 10 Hz stimulation (*n* = 5 preparations for each genotype, each the mean of 9 trials; see Methods for selection criteria). Trials were averaged together after eliminating the stimulus artifact before further analysis (see Supplementary Figure S1 for subtraction procedure). Synaptic strength was measured as the current integral of responses, then normalized by the mean baseline response. **(c)** Time course of changes in synaptic strength during the entire experiment. Traces at top are mean responses across all preparations during 10 Hz stimulation after eliminating stimulus artifacts and averaging across trials. **(d)** Replot of time course during 10 Hz stimulation only, on an expanded time scale. The 10 Hz train induced ∼5-fold enhancement at wildtype and SypFamilyKO synapses alike. **(e)** Replot of time course after 10 Hz stimulation only; both axes are expanded. The quickly decaying component – complete in less than 1 s – is termed facilitation and the slowly decaying component is termed augmentation [24, 46]. The theoretical curve is the single exponential *A e*^−^*^t/τ^* + 1 where *A* = 1.5 and *τ* = 7 s, which is characteristic of augmentation.

We induced facilitation with 10 Hz trains for the experiments in 0.5 mM Ca^2+^, rather than with pairs of pulses, because previous studies in primary cell culture and at neuromuscular junctions showed that facilitation continues to increase synaptic strength after the 2^nd^ pulse, which is an additional indication that lowering Ca^2+^ to 0.5 mM relieves occlusion [23, 24]. Nevertheless, paired pulse facilitation that is directly comparable to experiments under standard conditions could be extracted from the results by analyzing the responses to the first two pulses of the 10 Hz trains (gray box in 1d). The amount at SypFamilyKO synapses was equivalent to the amount at wildtype (i.e., 2.5 *±* 0.5-fold for SypFamilyKO vs 2.0 *±* 0.2-fold for wildtype; n = 5 for each). The result argues against concerns that the mechanism underlying induction of facilitation might have been deficient at SypFamilyKO synapses, and consequently strengthens the hypothesis that the deficit seen under standard conditions was caused by occlusion of the expression mechanism.

The overall increase in synaptic strength during the 10 Hz trains was substantially more than the paired pulse facilitation under standard conditions (compare Fig 1d to 1a). However, the larger amount was expected from previous studies in primary cell culture, and would be expected to grow even more with higher stimulation frequencies [24]. Similarly large amounts of facilitation are not seen during train stimulation under standard conditions [3], both because of limitations imposed by the ceiling value for *p_v_* noted in the Introduction, and because measurements of facilitation are confounded under standard conditions by simultaneous depletion of release sites [8].

### Deficit in long-lasting component of LTP

The previous evidence for a deficit in LTP pertained to synapses missing Syp and Syngr1 [5], and had not been confirmed for SypFamilyKO synapses, which are additionally missing Synpr and Syngr3. In experiments designed to confirm the deficit, we found that robust LTP was sustained for *>*100 min at wildtype synapses as expected, but ran down almost completely at SypFamilyKO synapses (compare white lines in Figs 2a and 2b, both are quantified in 2e). For these experiments, extracellular Ca^2+^ was 2 mM, matching the previous study, rather than the 2.6 mM used above for studying facilitation.

**Figure 2.**
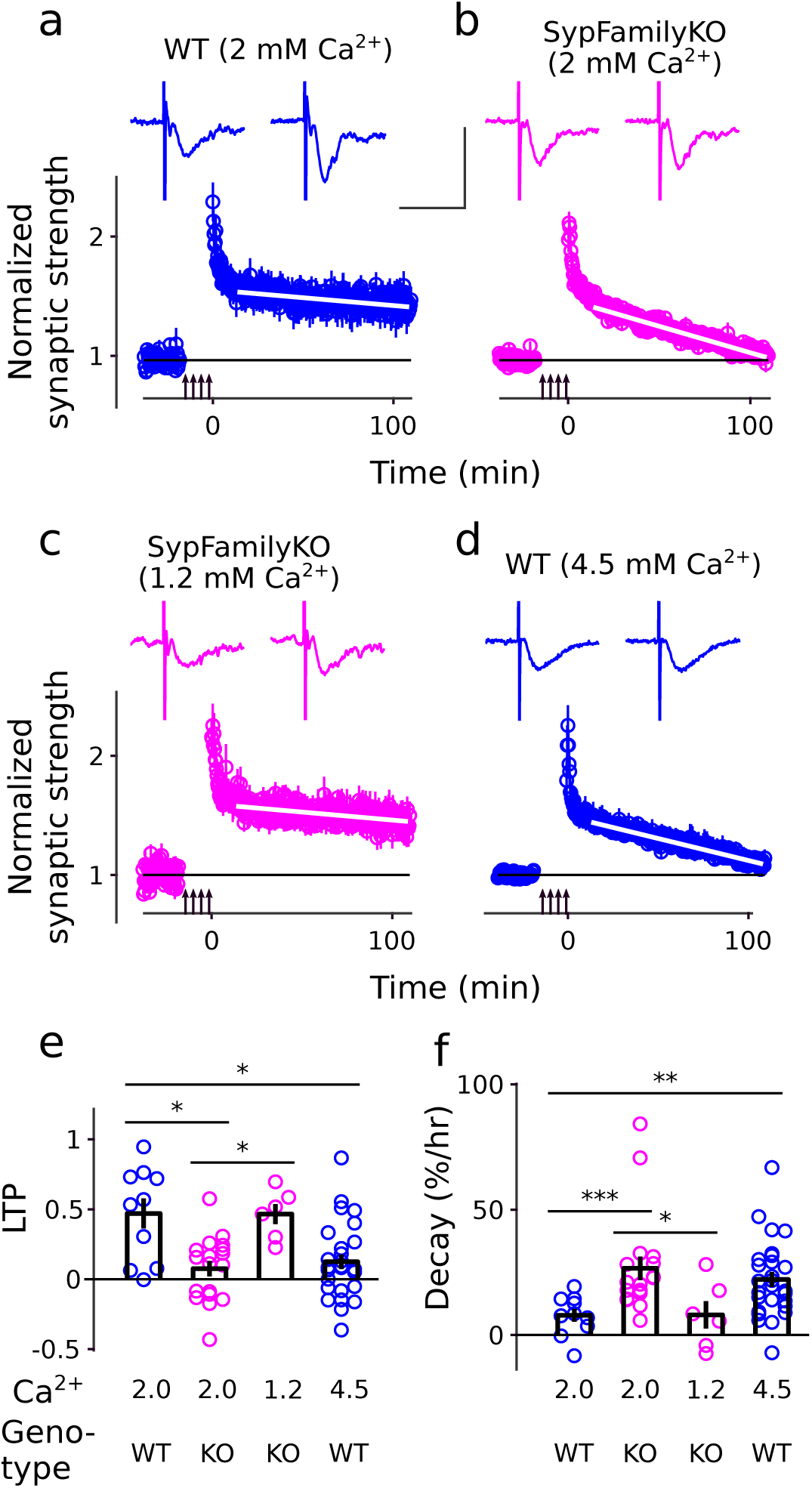
LTP after heavy induction: rescue in SypFamilyKO and recapitulation in wildtype. LTP induction was 4 trains of 100 Hz stimulation separated by 5 min intervals (vertical arrows). Mg^2+^ was 2 mM for all 4 experiments. Traces in insets are examples of recordings before and 100 min after induction; scale bars are 1 mV by 20 ms. **(a & b)** 2 mM Ca^2+^. Wildtype and SypFamilyKO experiments were interleaved, and the experimenter was blind to genotype. **(c & d)** SypFamilyKO experiments in 1.2 mM Ca^2+^ and wildtype in 4.5 mM Ca^2+^ were done as separate follow-on experiments, but the wildtype in 4.5 mM Ca^2+^ was interleaved with the SypFamilyKO experiments shown in Figure 3d. **(e)** LTP was quantified as the mean minus baseline for minutes 101 − 110 after induction. (* signifies *p <* 0.05, rank sum, *n* 6). Analysis was blind to genotype and Ca^2+^ level. **(f)** Decay was the slope between minute 15 and 110, matching the white lines in time course plots in Panels a-d (* signifies *p <* 0.05, ** *p <* 0.01, *** *p <* 0.001).

The result confirmed that there is indeed a major deficit in LTP at SypFamilyKO synapses. However, unlike here, LTP at knockout synapses in the previous study was already partly eliminated 10 min after induction, and did not run down further during the next 1 hr [5]. One methodological difference in addition to the difference in genotype is that we induced LTP with 4 trains of 100Hz stimulation, separated by 5 min intervals, rather than the 2 trains separated by 20 s used in the previous study. Nevertheless, LTP continued to run down at SypFamilyKO synapses after inducing with the 2 train protocol (Supplementary Figure S2a).

### Rescue of NMDA receptor-dependent LTP by lowering Ca*^2+^*

LTP at Schaffer collateral synapses is thought to be sustained by multiple maintenance mechanisms that differ in the time course of reversion to baseline [6]. The run-down at SypFamilyKO synapses initially seemed to suggest a deficit in the mechanism that maintains the longest lasting component. However, a viable alternative hypothesis that fits better with the new information about facilitation, above, was the longest lasting component of LTP is ordinarily expressed as an increase in *p_v_*, which is selectively occluded at SypFamilyKO synapses under standard conditions because the elevated baseline value prevents further increases. The continued expression of earlier components of LTP does not argue against occlusion of expression because the mechanisms underlying expression could be distinct for earlier components [6, 25].

We reasoned that the two hypotheses could be distinguished by attempting to measure LTP after lowering the extracellular Ca^2+^. The idea was that lowering Ca^2+^ would either have no effect, or would decrease LTP, if the deficit was at the level of maintenance. In contrast, like for facilitation, lowering Ca^2+^ would reverse any deficits at the level of expression. And indeed, as predicted by the occlusion hypothesis, the deficit in the longest lasting component of LTP was rescued by lowering extracellular Ca^2+^; in this case, from 2 to 1.2 mM (Fig 2c,e,f after inducing with 4 trains and Supplementary Figure S2b after inducing with 2 trains). And, lowering extracellular Ca^2+^ to 1.2 mM decreased baseline synaptic strength by a factor of 2.4 *±* 0.1 (n = 15) (Supplementary Figure S3a), which is more than the ∼2-fold elevation in baseline *p_v_* observed when comparing SypFamilyKO to wildtype [3]. Follow on experiments confirmed that lowering Ca^2+^ to 1.2 mM had no effect on the size of postsynaptic responses to spontaneous synaptic events at SypFamilyKO synapses (i.e., *minis*; Supplementary Figure S4), in-line with the long-standing premise that manipulating Ca^2+^ levels alters synaptic strength by altering *p_v_*, which is a presynaptic parameter [9, 11–13, 26]. The results are consistent with the occlusion hypothesis that the longest-lasting component of LTP is expressed via an increase in *p_v_* that is occluded by a ceiling value at SypFamilyKO synapses when Ca^2+^ is 2 mM, but not when 1.2 mM. However, the results do not completely rule out alternative explanations, which are assessed and discarded below.

### Recapitulation at wildtype rules out gain of function

For example, LTP at some other synapse types is, like facilitation, wholly presynaptic, and does not require NMDA receptor activation [27]. Because of this, we considered the possibility that key mechanisms underlying LTP at SypFamilyKO Schaffer collateral synapses might be fundamentally different compared to wildtype owing to a gain of function. However, all components of LTP continued to be blocked by NMDA receptor blocker APV at SypFamilyKO synapses, confirming that the LTP rescued at SypFamilyKO synapses by lowering extracellular Ca^2+^ requires NMDA receptors (Supplementary Figure S2b).

A related concern involved uncertainty about the amount of Ca^2+^ ordinarily needed to induce LTP. That is, induction requires influx of Ca^2+^ via NMDA receptors, and at least one previous study concluded that lowering extracellular Ca^2+^ to 1 mM reduced the influx below the threshold amount needed [28]. However, the finding that LTP could be induced at SypFamilyKO synapses when extracellular Ca^2+^ was 1.2 mM did not indicate that the induction mechanism was more sensitive to Ca^2+^ because equivalent LTP was induced under the same conditions at wildtype synapses (Supplementary Figure S5). A possible explanation is that the previous experiments were conducted without saturating amounts of NMDA receptor co-agonist glycine, the absence of which may have reduced Ca^2+^ influx by reducing receptor open probability. In any case, the continued presence of LTP at wildtype synapses in 1.2 mM Ca^2+^ fits with the widespread premise that LTP is involved in learning and memory because the Ca^2+^ concentration in vivo is likely close to 1.2 mM [29]. And indeed, no gross memory deficits were detected in SypFamilyKO mice in standard behavioral assays, including: novel object recognition; the Y-maze spontaneous alteration test; and a test for social memory (Supplementary Figure S6).

Finally, to rule out gain of function more comprehensively, we reasoned that it would be possible to recapitulate the deficit at wildtype synapses by elevating the baseline *p_v_* to SypFamilyKO levels. In experiments designed to test this, we found that increasing extracellular Ca^2+^ to 4.5 mM increased baseline synaptic strength by a factor of 2.3 *±* 0.15 (n = 21; Supplementary Figure S3b), which is at least as much as the ∼2-fold increase caused by knocking out synaptophysin family proteins [3]. And, the increase recapitulated the deficit in LTP (compare Fig 2b and 2d and bars 2 and 4 in Fig 2e). This result supports further the hypothesis that expression of the longest-lasting component of LTP is occluded at SypFamilyKO synapses by the elevated baseline *p_v_* by ruling out residual concerns that the mechanisms underlying LTP at SypFamilyKO synapses might be fundamentally different than at wildtype.

### Increasing *p_v_* further eliminates early component of LTP

Next, we discovered that the early component of LTP, within the first 30 min of induction, could also be eliminated by raising *p_v_* further. That is, the early component was not substantially altered at SypFamilyKO synapses under standard conditions (Fig 3a and 3b and bar 1 and 2 in Fig 3f), or at wildtype synapses in 4.5 mM Ca^2+^ (Fig 3c and bar 4 in 3f). However, the early component was nearly completely eliminated at SypFamilyKO synapses in 4.5 mM Ca^2+^ (Fig 3d and bar 3 in 3f). Follow-on experiments then showed that the deficit in the early component could be recapitulated at wildtype synapses, when Ca^2+^ was 4.5 mM, by decreasing Mg^2+^ from 2 to 0.5 mM (Fig 3e and bar 5 in 3f), which raises *p_v_* even further, likely by relieving partial block of voltage gated Ca^2+^ channels [9, 30]; we chose to raise baseline *p_v_* for these experiments by lowering Mg^2+^ to avoid increasing Ca^2+^ beyond the range routinely used in LTP experiments. Taken together, the results suggest that both early and long-lasting components of LTP can be eliminated by increasing the baseline *p_v_*, with earlier components eliminated at higher values of *p_v_*.

**Figure 3.**
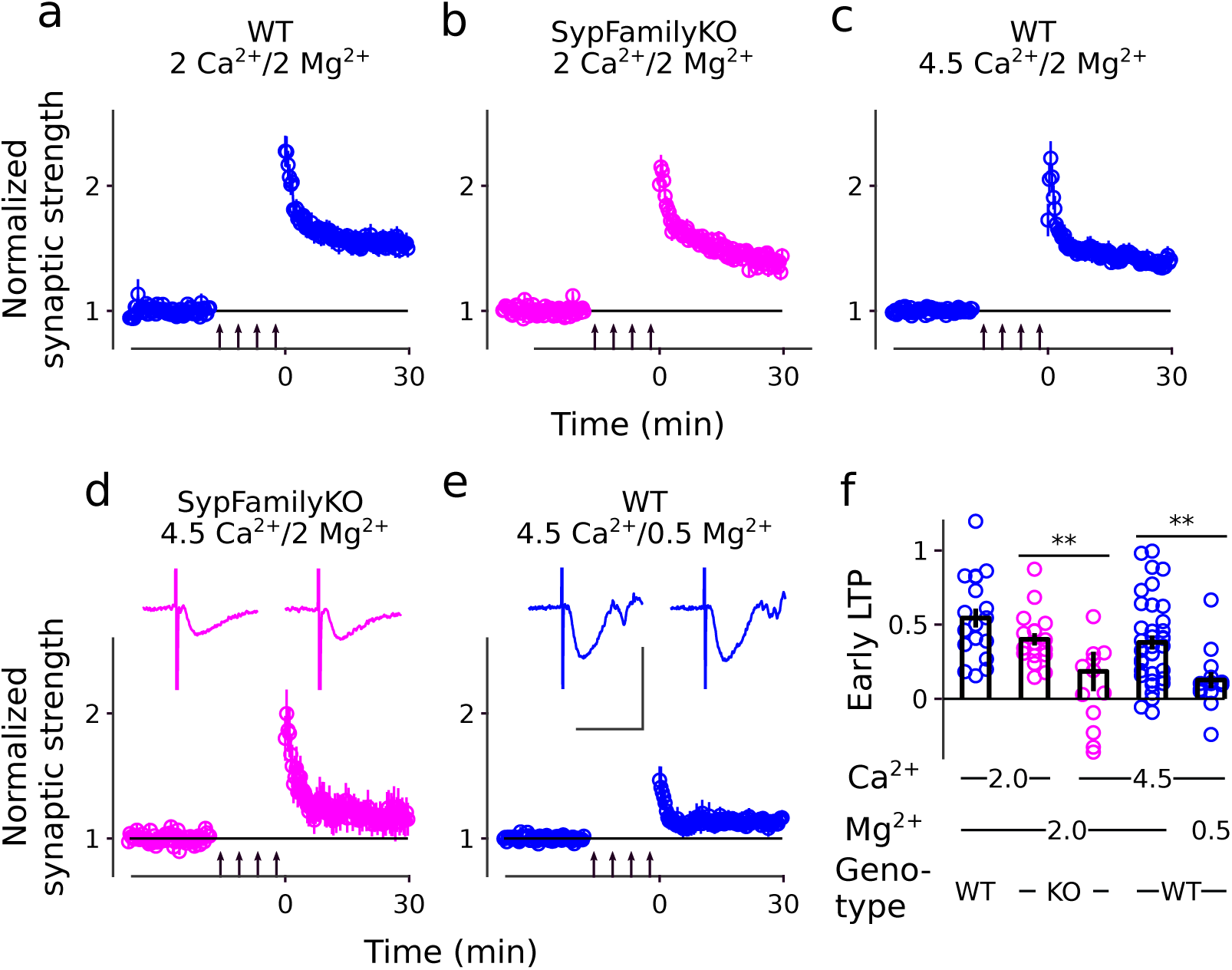
Early component of LTP is eliminated by raising p_v_ further. First 30 min after inducing LTP with 4 trains of 100 Hz stimulation for a range of Ca^2+^/Mg^2+^. **(a)** Includes experiments partly documented in Fig 2a. **(b)** Includes experiments partly documented in Fig 2b. **(c)** Includes experiments partly documented in Fig 2d. **(d)** Experiments were interleaved with experiments in **(c)**, and experimenter was blind to genotype. Scale bars are 1 mV by 20 ms. Independent further experiments with a similar result are documented in Fig 5a, left panel. **(e)** Includes experiments partly documented in Fig 4b. Scale for example traces is the same as in **(d)**. **(f)** Early LTP was measured between 25 and 30 min after stimulation (*n* 13; ** signifies *p <* 0.01, rank sum).

The conclusion is surprising given substantial evidence for addition of AMPA-type glutamate receptors to postsynaptic membranes within 20 – 40 min after LTP induction because new AMPA receptors would be expected to increase synaptic strength, but the increase would not be expected to be occluded by elevating baseline *p_v_* [14]. Because of this, we wondered if the elimination of the early component of LTP might have been caused by a general effect of the elevated baseline synaptic strength that was only indirectly related to the elevated *p_v_*. In experiments designed to test this, partially blocking AMPA receptors with 0.3 ➭M DNQX reduced synaptic strength by ∼ 60 %, meaning that synapses were _∼_ 2.5-fold stronger in the absence of DNQX. The difference was greater than the increase seen when switching Ca^2+^/ Mg^2+^ from 4.5/2 to 4.5/0.5 mM (Supplementary Figure S3c). Even so, the DNQX did not rescue the early component when Ca^2+^/ Mg^2+^ was 4.5/0.5 mM, or prevent the early component when 4.5/2.0 mM (Supplementary Figure S7). The combination of results indicates the deficit in early LTP when Ca^2+^/ Mg^2+^ was 4.5/0.5 mM must have been caused by the elevated baseline *p_v_* rather than by a general increase in synaptic strength. See Discussion for how the addition of postsynaptic AMPA receptors might fit with these findings.

### Intact induction

Next, to rule out the possibility that raising *p_v_* somehow interferes with induction of LTP, we found that the longest-lasting component could be unveiled at SypFamilyKO synapses after inducing in 2 mM Ca^2+^ by subsequently lowering Ca^2+^ to 1.2 mM. In contrast, no LTP was unveiled when omitting the 100 Hz stimulation used for induction (Fig 4a, controls were interleaved). Additional interleaved controls confirmed that the longest-lasting component of LTP was eliminated when Ca^2+^ was maintained at 2 mM (Supplementary Figure S8). These results support further the hypothesis that elevating baseline *p_v_* occludes the expression of LTP by demonstrating that the mechanism underlying induction is intact at SypFamilyKO synapses when extracellular Ca^2+^ is 2 mM.

**Figure 4.**
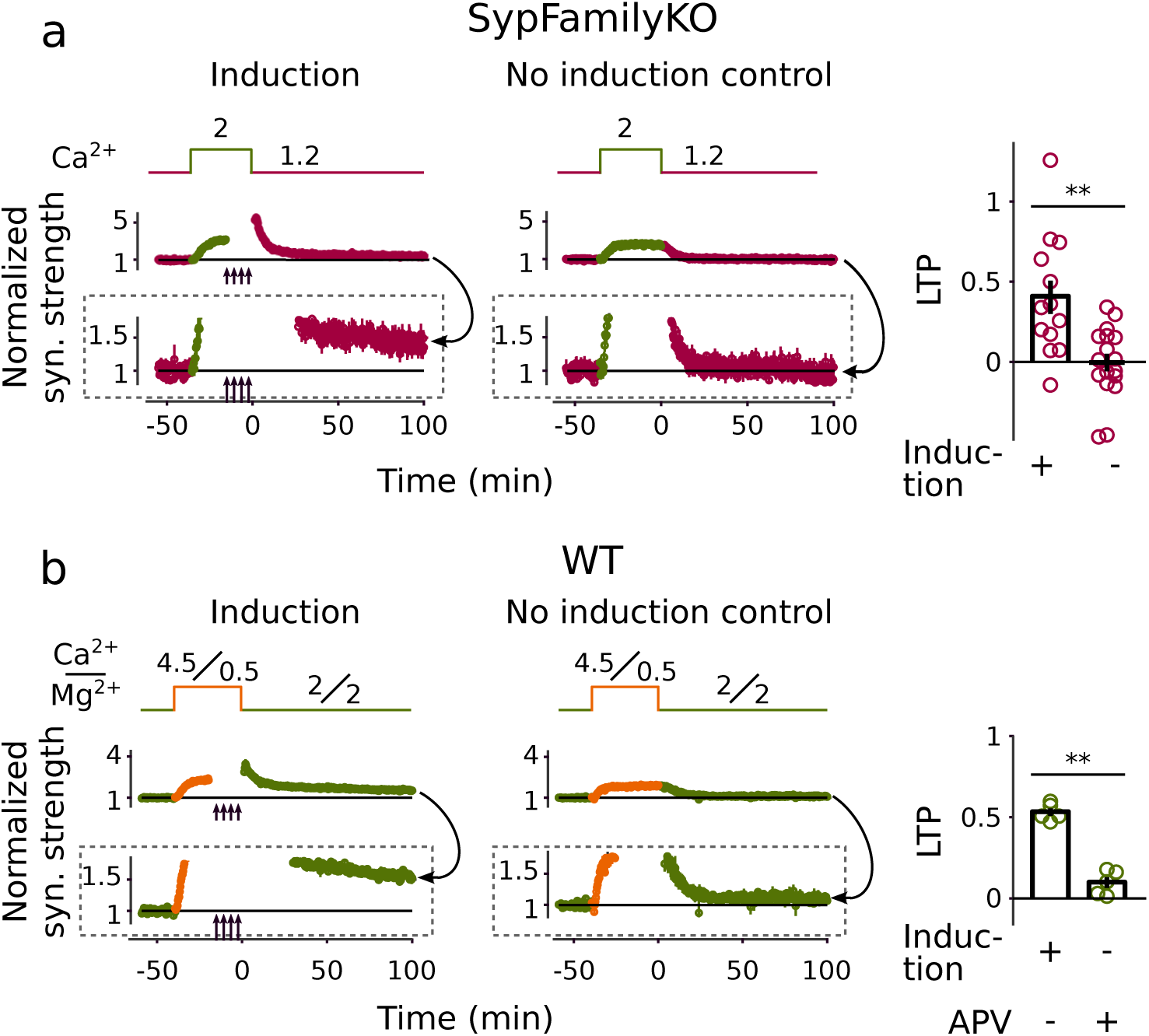
Intact LTP Induction. For each time course, the boxed lower plot is the same as the upper plot, except with expanded y-scale to highlight the presence or absence of LTP. LTP was quantified as the mean minus baseline for minutes 91 – 100 after induction. Baseline was established twice, first at the lower Ca^2+^/Mg^2+^, then again at the higher level. Time allowed to raise Ca^2+^ levels after the initial baseline varied a small amount between trials, but this is omitted from the time course plots. **(a)** SypFamilyKO. Trials with and without LTP induction were interleaved; ** signifies *p <* 0.01, rank sum, *n* 13. Mg^2+^ was 2 mM throughout all trials. Interleaved controls where Ca^2+^ was maintained at 2 mM after LTP induction are documented in Supplementary Figure S8. **(b)** Wildtype. 50 ➭M APV was present throughout trials with no induction; these were not interleaved, but interleaved trials with no induction and no APV are documented in Supplementary Figure S9, along with interleaved trials where Ca^2+^/Mg^2+^ was maintained at 4.5/0.5 mM throughout.

We obtained an analogous result for wildtype synapses that additionally pertained to the early component of LTP when Ca^2+^/Mg^2+^ was 4.5/0.5 mM during induction and 2/2 mM during measurement (Fig 4b). For these experiments, synaptic strength did not fully return to baseline in interleaved controls when we switched back to 2/2 mM without inducing LTP with 100 Hz stimulation (Supplementary Figure S9). However, the failure to fully return to baseline was likely caused by Ca^2+^ influx into the postsynaptic compartment via NMDA receptors that induced a minor amount of LTP during the low frequency (0.05 Hz) stimulation used to monitor synaptic strength, and not by a technical failure to fully replace the 4.5/0.5 mM solution with 2/2 mM (data not shown). And indeed, synaptic strength did fully return to baseline in follow-on experiments where NMDA receptor blocker APV was included throughout (Fig 4b). In any case, the potentiation in synaptic strength unveiled after switching to the 2/2 mM solution in the absence of APV was substantially greater when the induction protocol was included compared to when omitted, confirming that the majority of the potentiation was induced by the 100 Hz stimulation (Supplementary Figure S9). Finally, for these experiments, we additionally interleaved control trials designed to confirm that neither potentiation nor substantial depression/run-down occurred in the absence of 100 Hz stimulation when Ca^2+^/Mg^2+^ was maintained at 4.5/0.5 mM (Supplementary Figure S9).

Taken together, the results in this section show that blocking early and long-lasting components of LTP by either eliminating Syp family members or by altering extracellular Ca^2+^/Mg^2+^ does not prevent the induction of LTP. As a consequence, the block must occur either at the level of maintenance or expression.

### Intact maintenance

To distinguish between maintenance and expression, we: induced LTP at SypFamilyKO synapses in 4.5 mM Ca^2+^; then waited 15 min to verify that potentiation was minimal; and then lowered Ca^2+^ to 1.2 mM. To simplify the experimental design compared to Fig 4, we omitted prior baseline measurements in 1.2 mM Ca^2+^ and instead assessed LTP expression by comparing synaptic responses after lowering Ca^2+^ to 1.2 mM to the corresponding responses during matched trials with no 100 Hz stimulation (Fig 5a).

**Figure 5.**
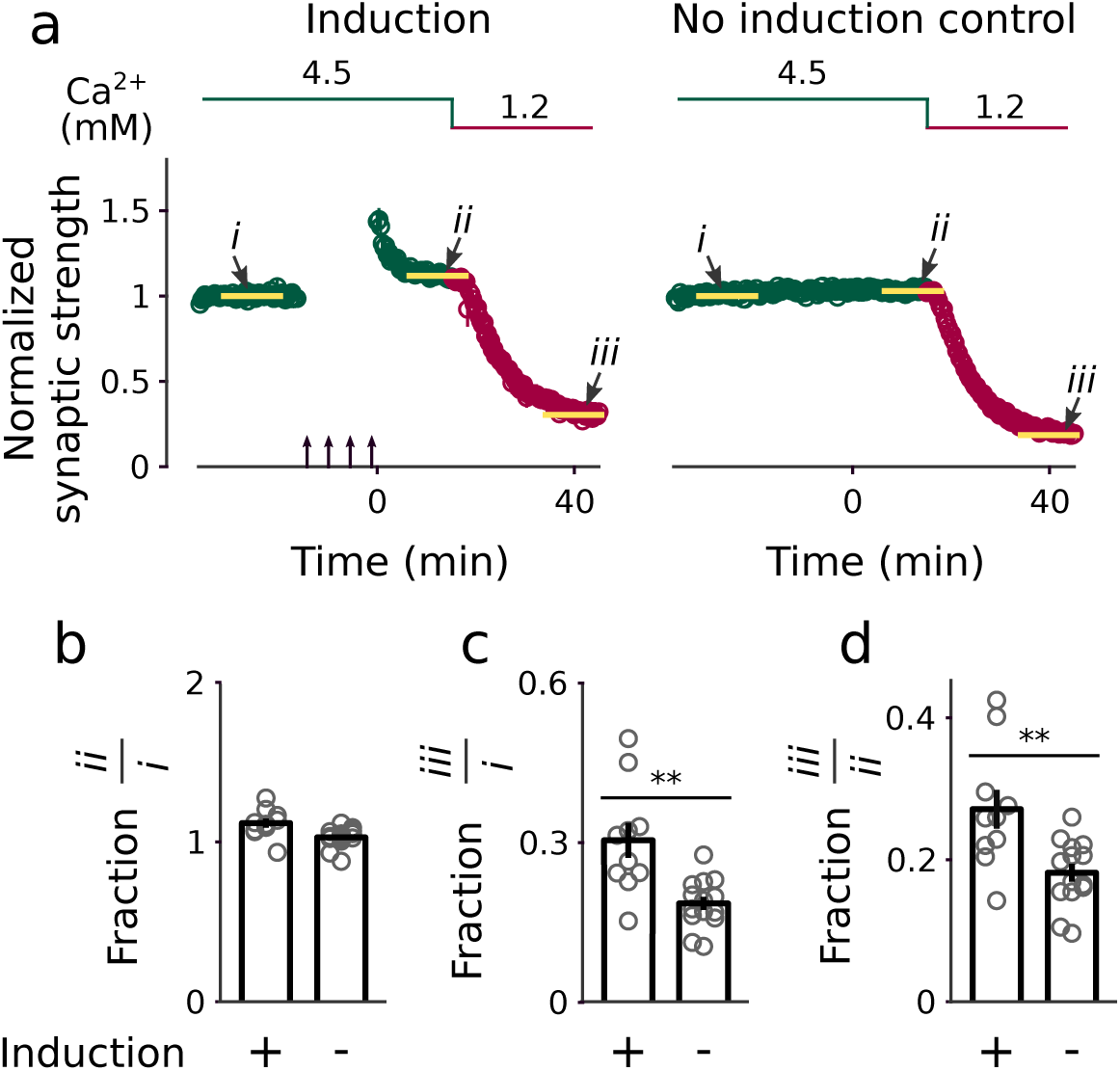
Intact LTP maintenance at SypFamilyKO synapses when Ca^2+^ is 4.5 mM. Induction and no induction control trials were interleaved. Induction was 4 trains of 100 Hz stimulation (upward arrows). Mg^2+^ was 2 mM throughout. **(a)** Downward arrows and yellow lines indicate windows that were averaged to quantify the results. **(b)** Synaptic strength 10 min after induction – which was when Ca^2+^ was still 4.5 mM – compared to baseline. **(c)** Synaptic strength when Ca^2+^ was 1.2 mM compared to baseline before LTP induction when Ca^2+^ was still 4.5 mM. Mean values were 0.30 *±* 0.04 (*n* = 10) for trials where LTP was induced and 0.19 *±* 0.01 (*n* = 14) for the no-induction controls. ** signifies *p <* 0.01 (rank sum). **(d)** Synaptic strength when Ca^2+^ was 1.2 mM compared to baseline 10 – 15 min after LTP induction when Ca^2+^ was 4.5 mM. Mean values were 0.27 *±* 0.03 for trials where LTP was induced and 0.18 *±* 0.01 for controls.

Potentiation had decreased to only 0.12 *±* 0.03 (*n* = 10) of baseline within 15 min after 100 Hz stimulation (Fig 5b), confirming the result, above, that LTP expression is minimal when extracellular Ca^2+^ is 4.5 mM. Nevertheless, the response remaining after lowering Ca^2+^ to 1.2 mM was 0.58-fold larger than the matched controls (Fig 5c), which is equivalent or more than the maximum amount of LTP induced under any conditions for either genotype above, or in comparable previous studies. The result shows that the absence of robust LTP expression when Ca^2+^ was 4.5 mM was not caused by interference with maintenance, and consequently confirms that the block occurs at the level of expression. As a consequence, the result, when taken together with the results in Figs 2 – 4, constitutes strong evidence for the hypothesis that, like facilitation, LTP expression is occluded by the elevated baseline *p_v_*.

A corollary is that LTP is manifest as a change in the relationship between extracellular Ca^2+^ and synaptic strength, and this was confirmed directly with additional analysis of the same experiments (Fig 5d).

## Discussion

Here we show that the mechanisms for maintenance and expression of at least two major components of LTP are separate at Schaffer collateral synapses and that expression of the two components can be blocked by elevating baseline *p_v_*, which is the efficiency with which action potentials trigger exocytosis of release ready synaptic vesicles within presynaptic terminals. Furthermore, the longest-lasting component could be blocked selectively, by elevating baseline *p_v_* to an intermediate level. Going forward, these findings provide an experimental methodology for dissecting LTP along two dimensions; maintenance versus expression, and early versus long-lasting components.

We manipulated baseline *p_v_* by both eliminating synaptophysin family proteins and by altering extracellular Ca^2+^/Mg^2+^. The two types of manipulations are orthogonal and effects were additive. That is: (a) Baseline *p_v_* is elevated at SypFamilyKO synapses. (b) Increasing Ca^2+^ from the physiological level of 1.2 mM up to 2 mM increases *p_v_* further and blocked a long-lasting component of LTP, whereas (c) it took a larger increase, to 4.5 mM, to recapitulate the result, at wildtype synapses. Finally, (d) increasing Ca^2+^ at SypFamilyKO synapses to 4.5 mM additionally blocked the early component of LTP, whereas (e) at wildtype synapses it was additionally necessary to lower Mg^2+^, which increases *p_v_* further by relieving Mg^2+^-dependent block of Ca^2+^ channels.

We cannot rule out still unidentified deficits at SypFamilyKO synapses that are not related to the elevated baseline *p_v_* [31], including subtle deficits in postsynaptic mechanisms [32]. However, manipulating extracellular Ca^2+^ and Mg^2+^ has long been recognized as a potent and quickly reversible method for altering *p_v_* [33], with no indication of confounding interference with postsynaptic mechanisms that could account for: the rescue of LTP expression at SypFamilyKO synapses; the recapitulation of the deficits in both early and long lasting components at wildtype; and the absence of interference with LTP induction or maintenance. In addition, a previous study showed that NMDAR-dependent LTP expression is blocked at neonatal Schaffer collateral synapses under standard experimental conditions, where baseline *p_v_* is elevated compared to mature synapses, and, like at mature SypFamilyKO synapses, here, can be rescued by lowering the extracellular Ca^2+^.

Future studies needing to isolate LTP expression from induction and maintenance, or the long-lasting component from earlier components, will not necessarily require SypFamilyKO mice because expression and the longest lasting component could both be isolated at wildtype synapses simply by manipulating Ca^2+^ and Mg^2+^. In any case, synaptophysin family proteins clearly are not essential for LTP despite severely reduced LTP expression in knockouts when Ca^2+^ was 2 mM or higher.

The most likely explanation for the blocked LTP expression when baseline *p_v_* is elevated seems to be that both early and long-lasting components of LTP are expressed via increases in *p_v_*, which are occluded when the baseline is already elevated. Indeed, a previous study, unrelated to LTP or SypFamilyKOs, showed that *p_v_* is highly susceptible to modulation by multiple factors that can occlude each other [8]. If so, occlusion would have to be multi-factorial to explain why the late and early components are occluded at distinct Ca*_2+_*/Mg^2+^ levels. The identity of the factors is not known, but our working model of presynaptic function, developed previously to explain rate-limiting steps in synaptic vesicle trafficking [34], already contains multiple types of release sites with distinct values for *p_v_* (Fig 6) that would be occluded at distinct Ca^2+^/Mg^2+^ levels. One possibility is that the longest lasting component of LTP is expressed by potentiating *efficient* release sites with inherently high *p_v_*, whereas the early component is expressed by potentiating *inefficient* release sites with low *p_v_*. An analogous mechanism was proposed previously to explain inconsistent evidence across studies for interactions between LTP and facilitation [35, 36].

**Figure 6.**
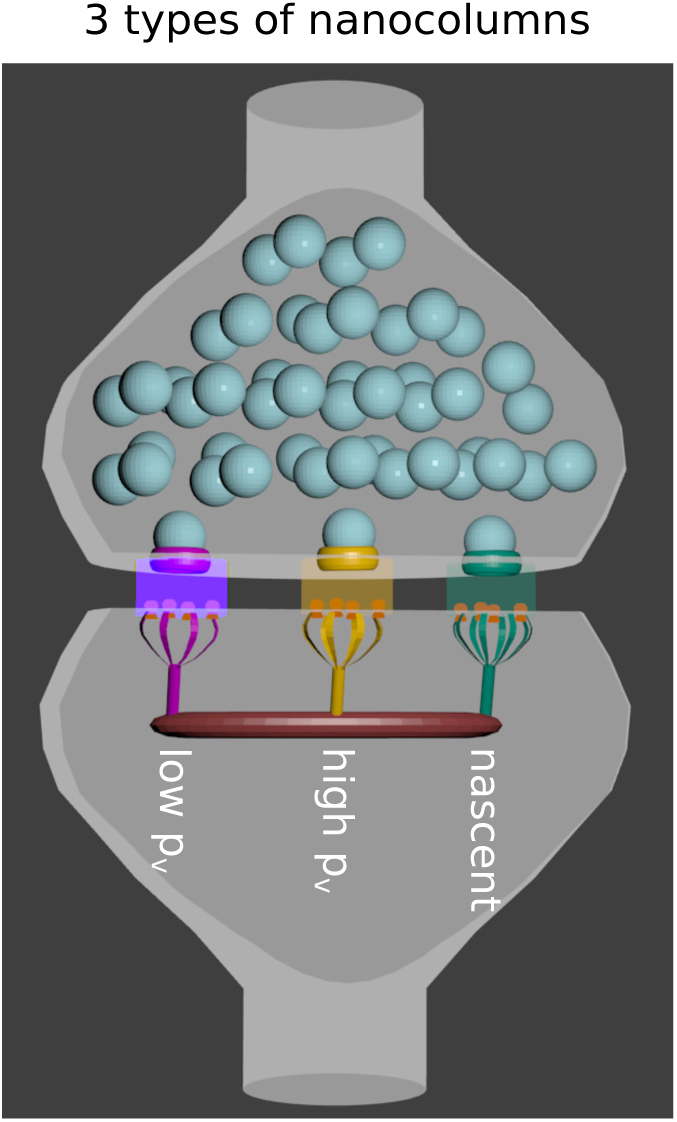
Working model. High and low *p_v_* nanocolumns contain release sites that are *efficient* and *inefficient*, respectively, at catalyzing exocytosis; these explain normal and reluctant readily releasable vesicles [3, 13, 34]. *Nascent* release sites are not functional during the first 2 hr after inducing LTP, and were added expressly to explain previous evidence for newly added release sites and postsynaptic receptors that do not seem to play a large role in LTP in the present study.

### Nanocolumn structural hypothesis

*p_v_* is by definition a presynaptic parameter, and the new support for a presynaptic LTP expression mechanism complements previous studies showing: (a) more neurotransmitter release [15]; (b) more exocytosis from presynaptic terminals measured with FM dyes [18]; and, (c) increases in probability of release at individual synapses measured with electrophysiological and optical imaging techniques [19, 22, 37]; along with (d) mathematical analyses of changes in trial-to-trial fluctuations in synaptic responses [16, 17]. However, *p_v_* pertains to individual release sites, and does not seem to be influenced by the number of sites or number of readily releasable vesicles [3, 12, 13]; indeed, the ceiling value for *p_v_* continues to limit facilitation at wildtype synapses when most release sites are unoccupied [8]. Because of this, our results do not directly support the fundamentally different presynaptic hypothesis that LTP is expressed by addition of new release sites that engender a larger readily releasable pool, despite strong evidence that inducing LTP increases the number of morphologically docked vesicles seen with electron microscopy [38].

One possibility, suggested previously, is that the newly docked vesicles might be docked to *nascent* release sites that are not yet functional during the first 2 hr after inducing LTP [38]. If so, the newly docked vesicles would not participate in synaptic transmission, and would not contribute to LTP during the time course of our experiments. Instead, the newly docked vesicles might participate in memory consolidation hours later. Alternatively, the newly added vesicles could be docked to the inefficient/low *p_v_* release sites mentioned above, which would not contribute much to synaptic transmission during the low frequency stimulation used to monitor LTP expression; inefficient/low *p_v_* release sites are thought to operate as high pass/low cut frequency filters [13] so new additions might selectively store information encoded by spike timing within bursts of action potentials [39].

In either case, the emerging concept of *transynaptic nanocolumns* where synaptic vesicle docking sites are linked across the synaptic cleft to dedicated clusters of postsynaptic neurotransmitter receptors [40] could then explain extensive evidence for increases in AMPA receptors within postsynaptic membranes [14, 41]. That is, the new nascent or inefficient/low *p_v_* release sites might be components of newly added nanocolumns that would include new clusters of receptors on the postsynaptic side of the synaptic cleft (Fig 6). The new receptors could be functional right away, but would not contribute much to LTP as measured with standard techniques because they would not participate in synaptic transmission during low frequency stimulation. And indeed, there is evidence that receptors added after the induction of LTP are added as part of new nanocolumns rather than to previously existing receptor clusters [42].

The hypothesis depends on the premise that neurotransmitter spillover between nanocolumns is minimal in ex vivo tissue. The premise has not been verified directly, but is consistent with evidence that postsynaptic responses generated by exocytosis of single vesicles at large Schaffer collateral synapses, with many release sites, are not larger than responses at small synapses even though the postsynaptic membranes likely contain fewer receptors [43, 44].

## Methods

SypFamilyKO and matched wildtype mice were from the colony described in ref. [3]. All electrophysiology experiments were conducted on 400 µm thick *ex vivo* transverse slices from hippocampus with CA3 removed as described in ref. [12] except slices were incubated at 34 C for 30-60 min immediately after slicing and transferring to an extracellular incubation/recording solution containing (in mM): 120 NaCl; 1.25 NaH*_2_*PO*_4_*; 26 NaHCO*_3_*; 3.5 KCl; 10 glucose; 0.05 picrotoxin; and CaCl*_2_* and MgCl*_2_* as indicated below. Synaptic responses were evoked with a Ag/AgCl*_2_* electrode inserted into a glass pipette with tip diameter 5 ➭m, filled with extracellular solution or 150 mM NaCl, and placed within *stratum radiatum*, about three-quarters of the distance between the principle neuron layer and *stratum lacunosum-moleculare*. Stimulation was biphasic with pulses lasting 100 ➭s and, in most cases, intensity 100 ➭A; pulses were constant voltage instead of constant current for some experiments.

**Short-term enhancement in low Ca^2+^:** Slices were from 13 – 21 day old mice (either sex). CaCl_2_ was 2.6 mM and MgCl_2_ was 1.3 mM in the incubation solution and in the extracellular recording solution at the start of experiments. Synaptic responses were recorded in whole cell voltage clamp mode by patch-clamping CA1 principle neurons. Intracellular recording solution contained (in mM): 125 KGluconate; 8 NaCl; 10 Hepes; 1 EGTA; 10 KCl; 0.2 CaCl_2_; 4 MgATP; 0.3 LiGTP, adjusted to pH 7.4 by adding KOH. Stimulus intensity and micro-electrode placement were configured before lowering Ca^2+^/Mg^2+^ to 0.5/1.5 mM, which typically lowered response size to a point where some trials resulted in synaptic transmission failures. Stimulation artifacts were eliminated by subtracting the mean failure from all responses (see Supplementary Figure S1). Trials were repeated multiple times, and were averaged together after eliminating the stimulation artifact, but before further analysis.

**Criteria:** Results from individual preparations were only included in the final analysis if the coefficient of variation of the 5 baseline responses after averaging across trials was less than 0.7, which eliminated preparations with excessive response-to-response scatter. Excessive scatter occurred for two reasons: (a) when few trials were available for averaging; and, (b), when the baseline probability of release was low. Stimulation was not limited to single afferents, so baseline probability of release was the sum of synapses from an unknown number of afferents; the number likely varied between experiments and would be expected to be more for wildtype that for SypFamilyKO slices, although the number was not estimated.

**LTP:** Slices were from 6 – 8 week old males. CaCl_2_ and MgCl_2_ for the incubation solution were both 2 mM and as indicated in the Results section for the recording solution. The recording solution additionally contained 20 ➭M glycine. Synaptic responses were recorded as field potentials in *stratum radiatum*, at approximately the same distance from *stratum lacunosum-moleculare* as the stimulating electrode. Up to 4 field potential recordings were conducted simultaneously on 2 electrophysiology workstations; each recording chamber held 2 independent slices. Experiments comparing stimulation protocols were typically interleaved, as indicated in Results and Figure Legends, but we did not conduct two-pathway experiments on the same slice because of possible interference between pathways of latent components of LTP [45]. Field potentials were measured as the slope of the rising phase of the synaptic response between 15 and 85 % of peak. **Criteria:** Results from individual preparations were only included if the baseline synaptic response did not change more than 6 % over 20 min of baseline when judged by linear regression, and if the stimulus artifact did not change perceptibly.

## Declarations

Funding This work was funded by: the Generalitat Valenciana Prometeo Excellence Program (CIPROM2022/8) and the Ministry of Science of Spain (BFU2016-80918-R, PID2019-111131GB-I00, and PID2023-153133NB-I00). The funders had no role in study design, data collection and analysis, or preparation of the manuscript.

Ethics approval All procedures were conducted in accordance with the European and Spanish regulations (2010/63/UE; RD 53/2013) and were approved by the Ethical Committee of the Generalitat Valenciana (2022/VSC/PEA/0255).

Authors’ contributions: SdO and JJRG conducted electrophysiological experiments and contributed to writing. JFW designed electrophysiological experiments, analyzed the data, and wrote the manuscript. RV designed, conducted and analyzed the behavioral experiments and contributed to writing.

## Acknowledgments

We thank: Dr. Sandra Jurado for teaching us the basics of LTP experiments; Drs. José Esteban and Robert Renden for helpful feedback during the evolution of this project; and Drs. Felix Leroy, Joaquin Piriz, and Isabel Pérez-Otaño for suggestions during writing.

---

**Supplementary Figure S1.**
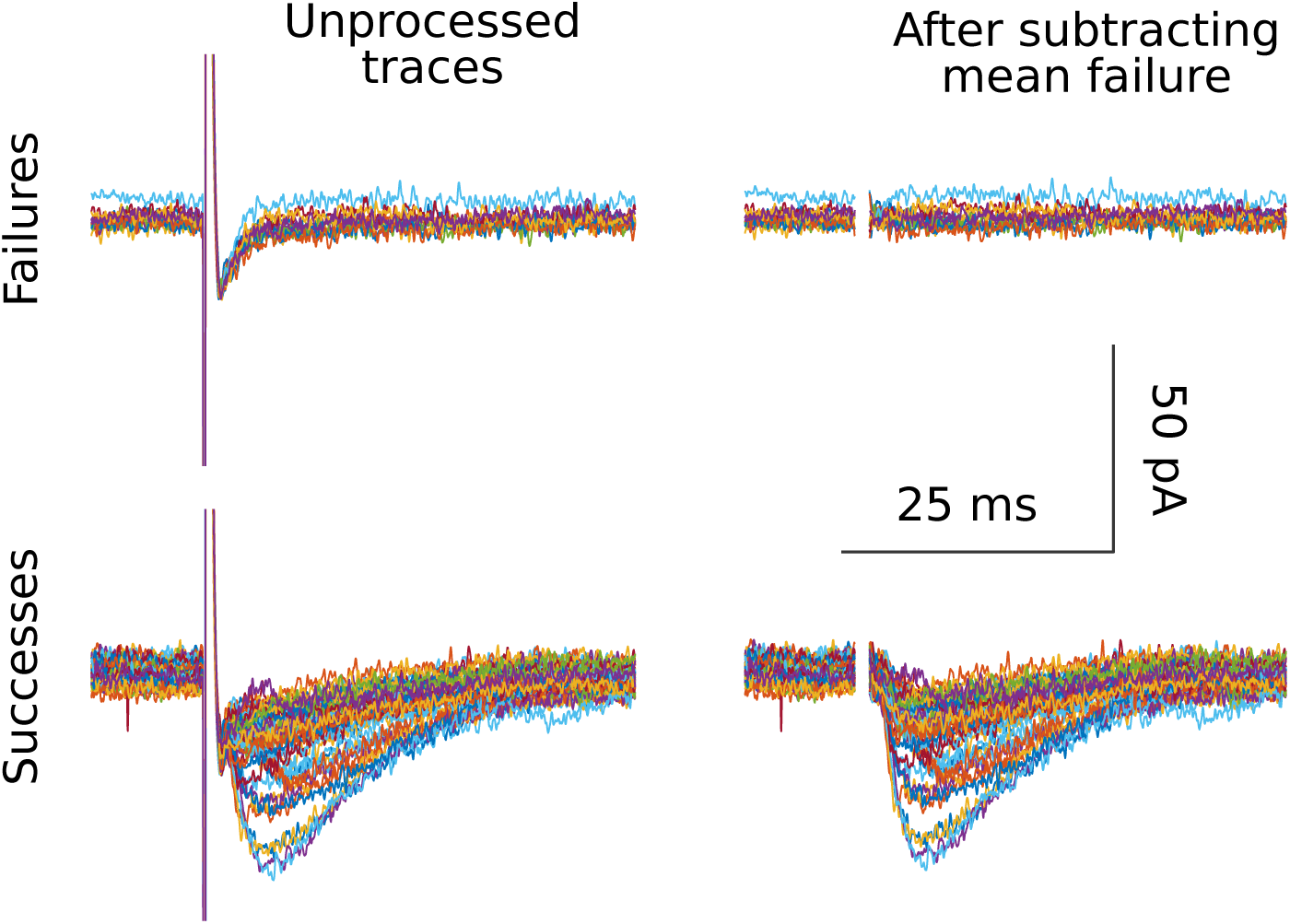
Procedure for eliminating stimulus artifacts in low Ca^2+^. Leftmost traces are responses judged to be *Failures* (upper) and *Successes* (lower). Rightmost traces are after digitally subtracting the mean failure. The example is from a SypFamilyKO preparation.

**Supplementary Figure S2.**
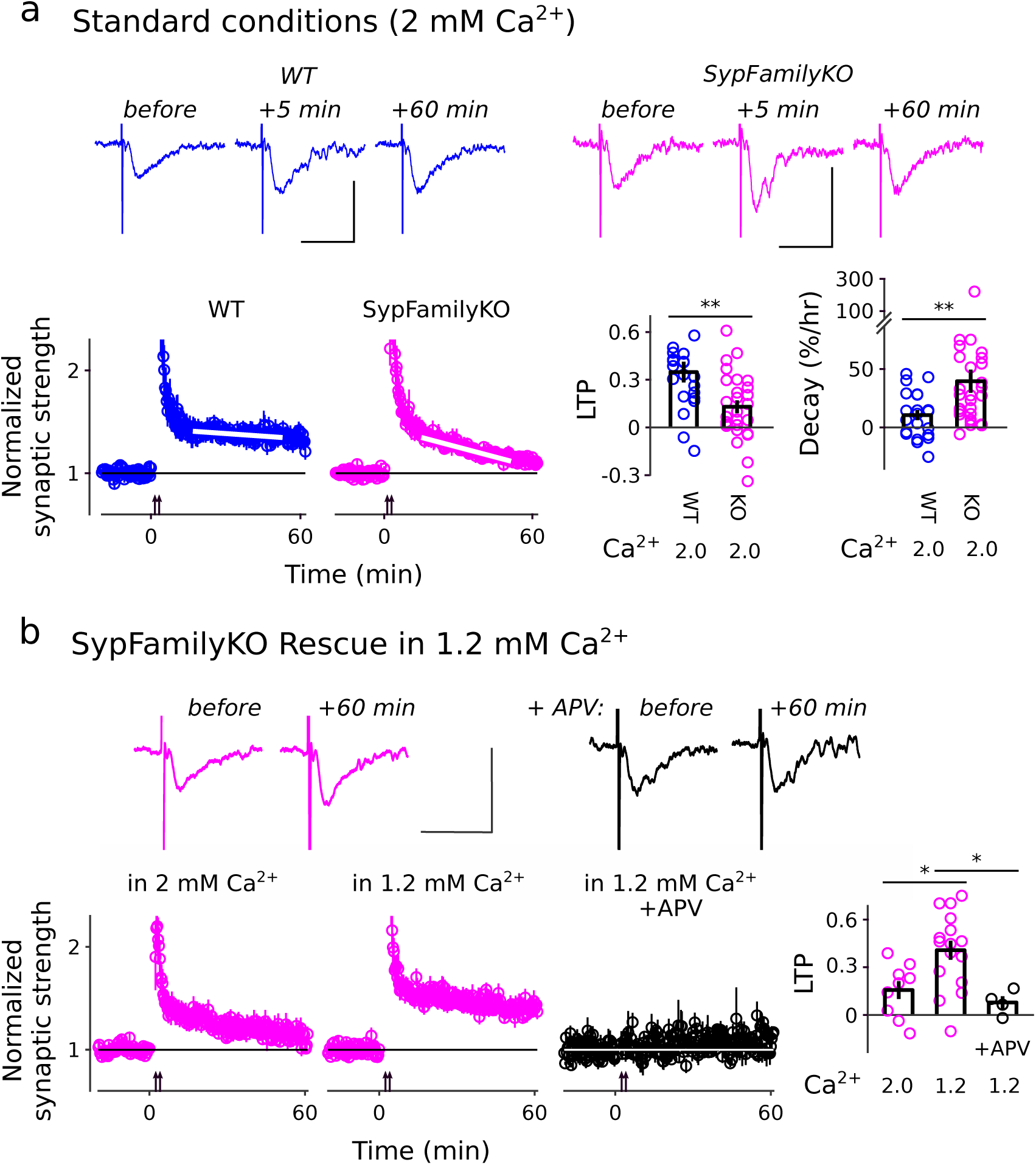
LTP induced by 21 s-long trains of 100 Hz *stimulation*. **(a)** Longest lasting component of LTP is eliminated at SypFamilyKO synapses under standard conditions when Ca^2+^/Mg^2+^ was 2/2 mM. Genotypes were interleaved and experiments and analysis were blind to genotype. Insets are example traces before and after attempting to induce LTP. Scale bars are 0.5 mV by 20 ms. y-axes of time course plots are truncated. For the bar graphs, LTP was quantified as the mean minus baseline for minute For the bar graphs, LTP was quantified as the mean minus baseline for minute 51 – 60 after induction and decay between minute 15 and 50. ** signifies *p <* 0.01, rank sum; *n* ≥ 21. **(b)** Rescue when Ca^2+^ is 1.2 mM. Experiments in 2 and 1.2 mM Ca^2+^ were interleaved. y-axes of time course plots are truncated. * signifies *p <* 0.05, rank sum; *n ≥* 4.

**Supplementary Figure S3.**
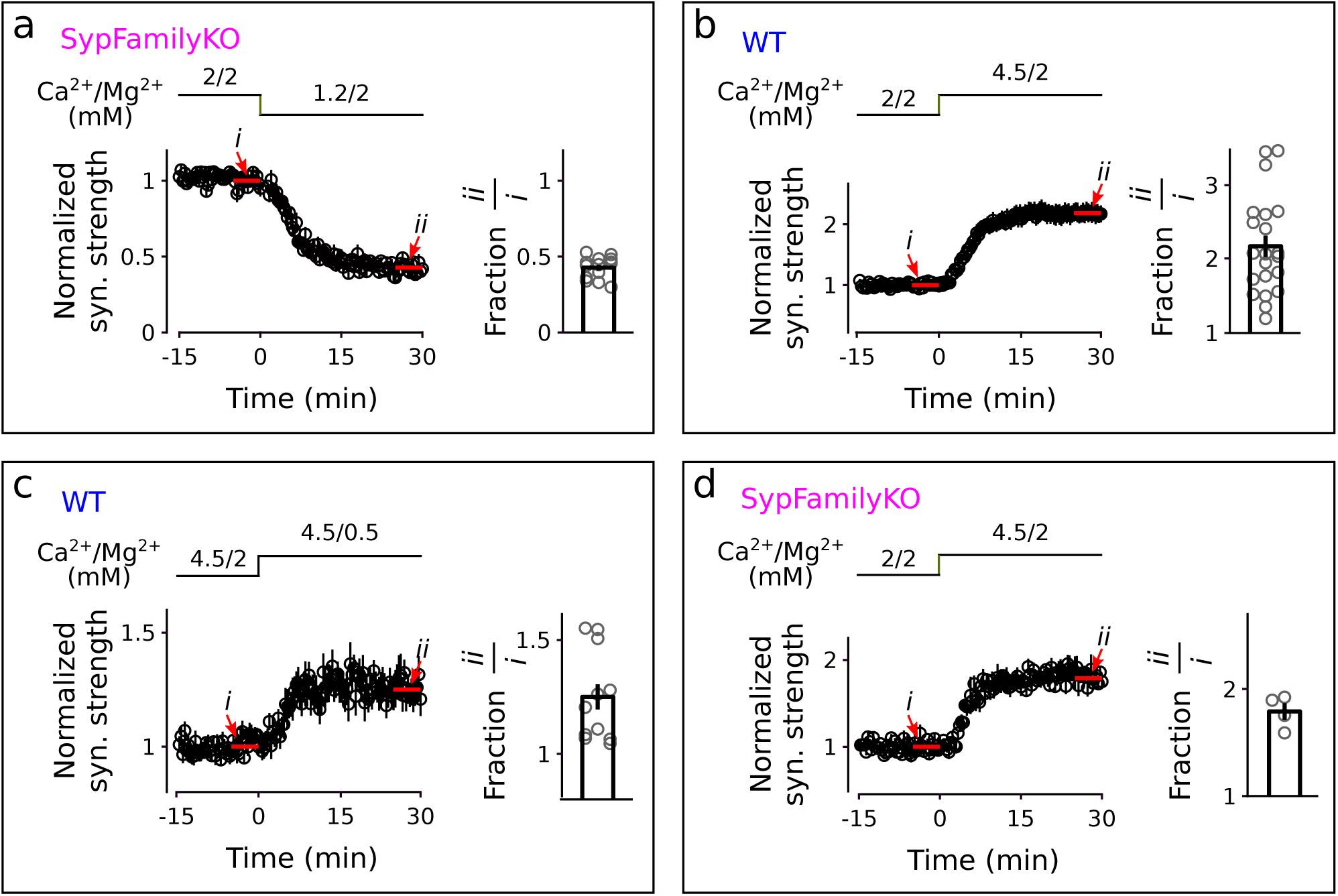
Ca^2+^/Mg^2+^ dependent changes in synaptic strength. **(a)** Responses are re-normalized from Figure 4a. Switching from 2 to 1.2 mM Ca^2+^ in 2 mM Mg^2+^ at SypFamilyKO synapses decreased synaptic strength to 0.43 *±* 0.02 (n = 15) of baseline, which was by a factor of 2.4 *±* 0.1 (i.e., calculated as the mean of the reciprocal values). Synaptic strength values used to calculate the fraction by comparing points *i* and *ii* are the mean of 10. **(b)** Switching from 2 to 4.5 mM Ca^2+^ in 2 mM Mg^2+^ at wildtype synapses increased synaptic strength by a factor of 2.3 *±* 0.15 (n = 21). **(c)** Switching from 2 to 0.5 mM Mg^2+^ in 4.5 mM Ca^2+^ increased synaptic strength at wildtype synapses by a factor of 1.25 *±* 0.06 (n = 12). NMDA receptors were blocked throughout with 50 ➭M APV. **(d)** Switching from 2 to 4.5 mM Ca^2+^ in 2 mM Mg^2+^ at SypFamilyKO synapses increased synaptic strength by a factor of 1.8 *±* 0.1 (n = 4). NMDA receptors were blocked throughout with 50 ➭M APV.

**Supplementary Figure S4.**
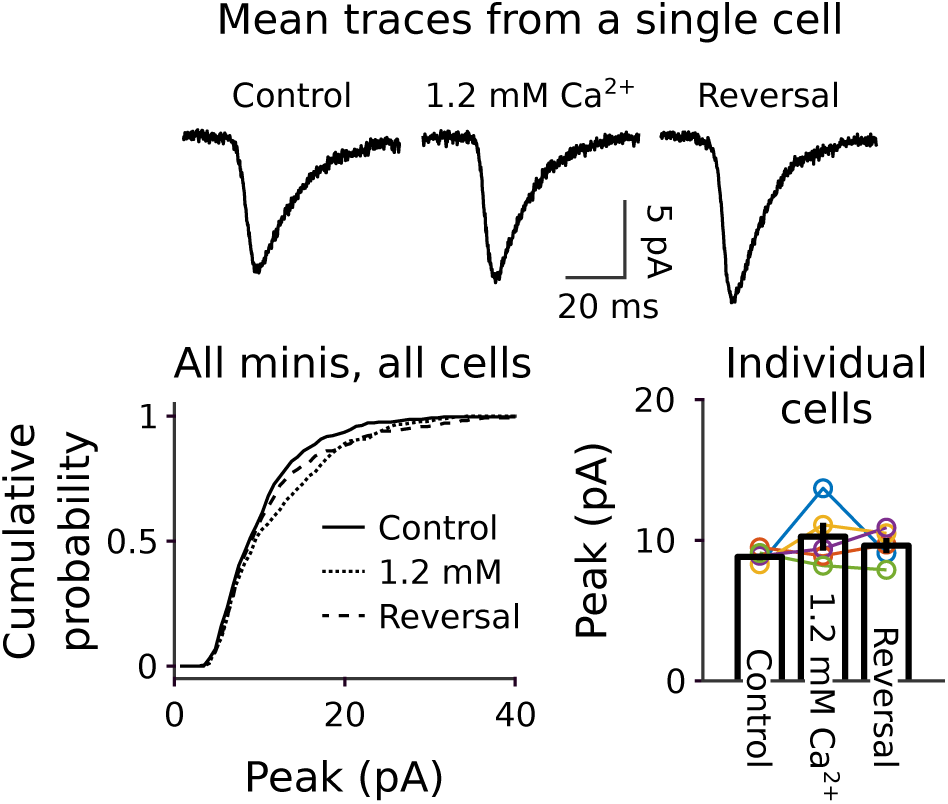
No change in size of minis when lowering Ca^2+^ to 1.2 mM at SypFamilyKO synapses. Minis were recorded in whole cell voltage clamp during 5 min intervals, first when extracellular Ca^2+^ was 2 mM, then after changing to 1.2 mM, and finally a third time after returning to 2 mM. Experiments were excluded if the access resistance changed more than 10 % (*n* = 5 neurons). **(a)** Mean electrophysiological traces for a single neuron. **(b)** Cumulative probability versus peak amplitude for all minis from all neurons. **(c)** Mean peak size of all minis from the individual neurons.

**Supplementary Figure S5.**
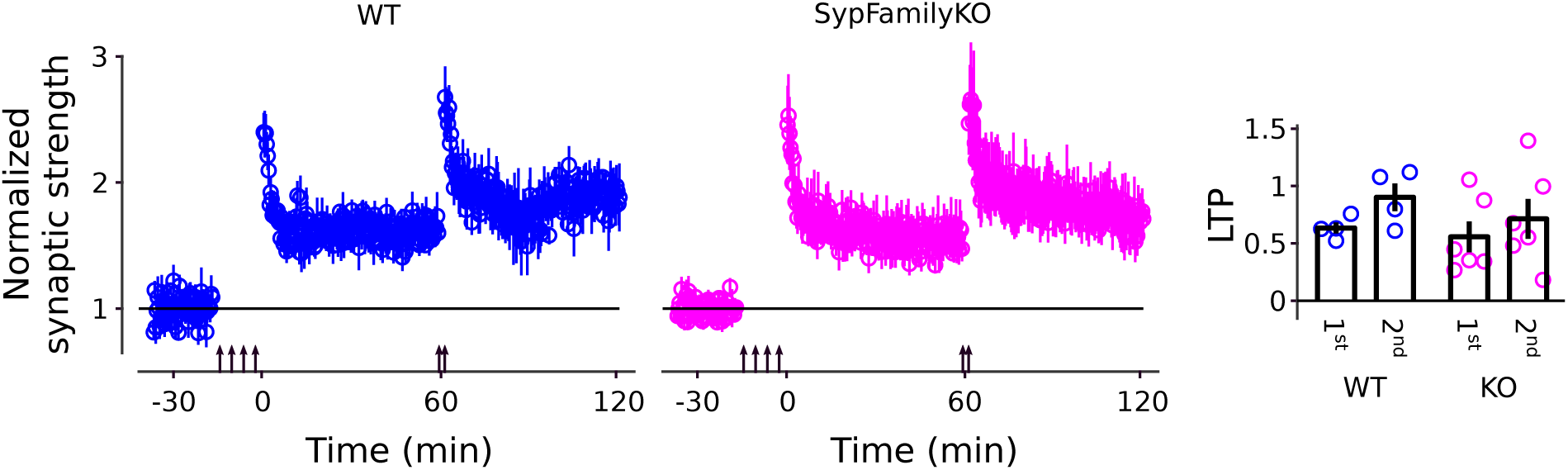
Comparison of LTP at wildtype and SypFamilyKO synapses in 1.2 mM Ca^2+^. LTP was first induced with 4 trains of 100 Hz stimulation as in Fig 2 and then, after 60 min, with the 2 train protocol as in Supplementary Figure S2. LTP was quantified as the mean minus baseline for minutes 51 – 60 after the 1^st^ induction protocol and for minutes 111 – 120 for the 2^nd^ (*n ≥* 4).

**Supplementary Figure S6.**
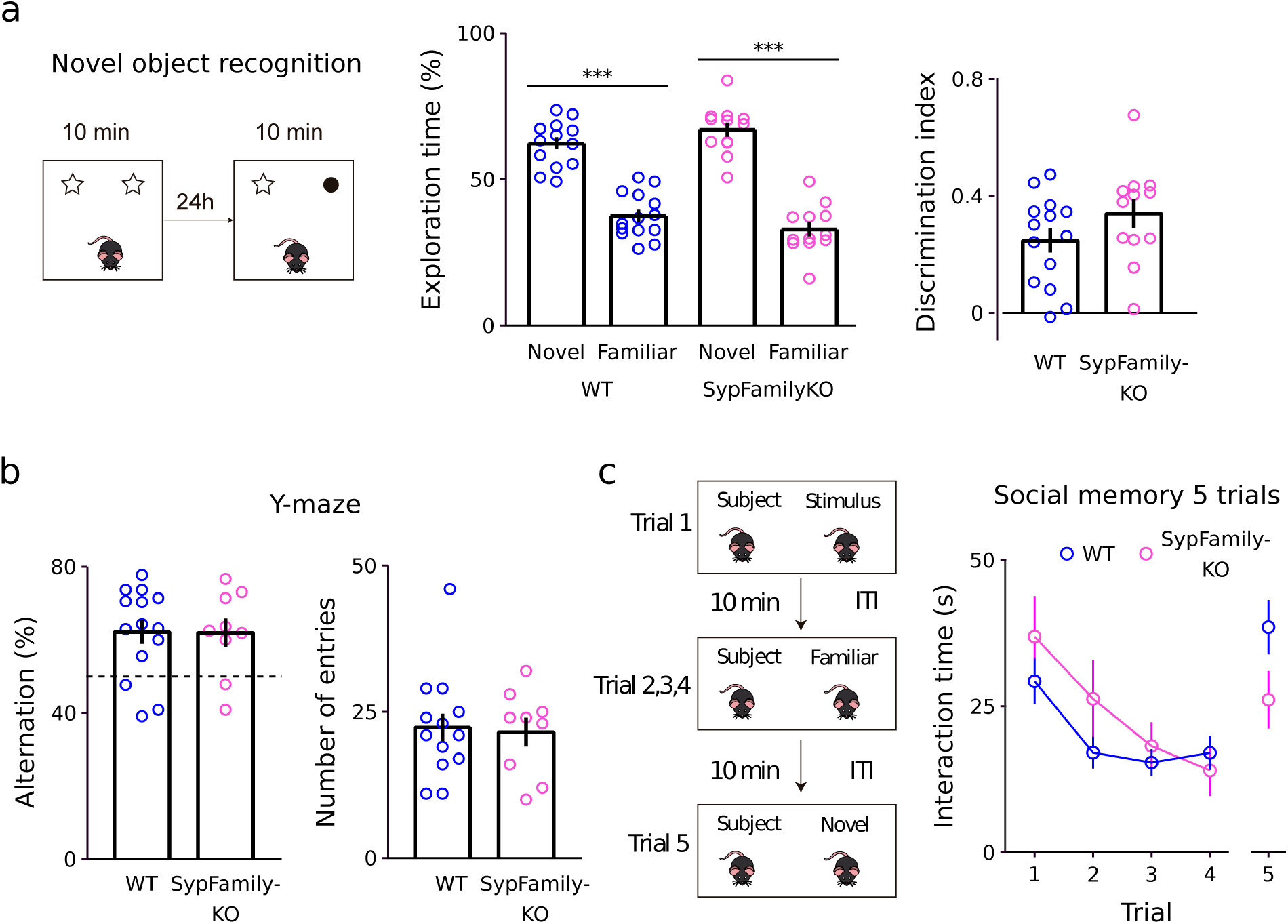
No obvious deficits in standard memory tasks. Experiments were recorded with a video camera and later scored by hand, blind to genotype. **(a)** Novel object recognition: Mice were first habituated to a 30 cm X 30 cm open arena for 10 min on day 1, then returned for 10 min on day 2 after adding 2 identical objects, and finally returned for 10 min on day 3 after replacing one of the objects with a novel object. Exploration time was total time when the mouse’s nose was within 2 – 3 cm of the object and pointed towards the object. Discrimination index is difference in time with novel and familiar object, normalized by total time with either object. *** signifies *p <* 0.0001, two-way ANOVA followed by Tukey’s multiple comparisons test. **(b)** Y-maze spontaneous alternation: Mice were placed in a Y-shaped maze with 3-arms and monitored for 8 min. Correct triads were noted when all 3 arms were entered sequentially. Alternation index is number of correct triads normalized by number of possible triads. Dashed line is chance (50 %). **(c)** Social memory 5 trials: Test mice were first habituated to a clean cage with dimensions matching the home cage for 30 min. A novel juvenile with matched sex was then introduced for 2 min intervals then removed for 10 min intervals for 4 trials. Finally, a second sex matched novel juvenile was introduced for the 5^th^ trial. Social interaction time was calculated as total time spent sniffing any part of the body, allogrooming and close following.

**Supplementary Figure S7.**
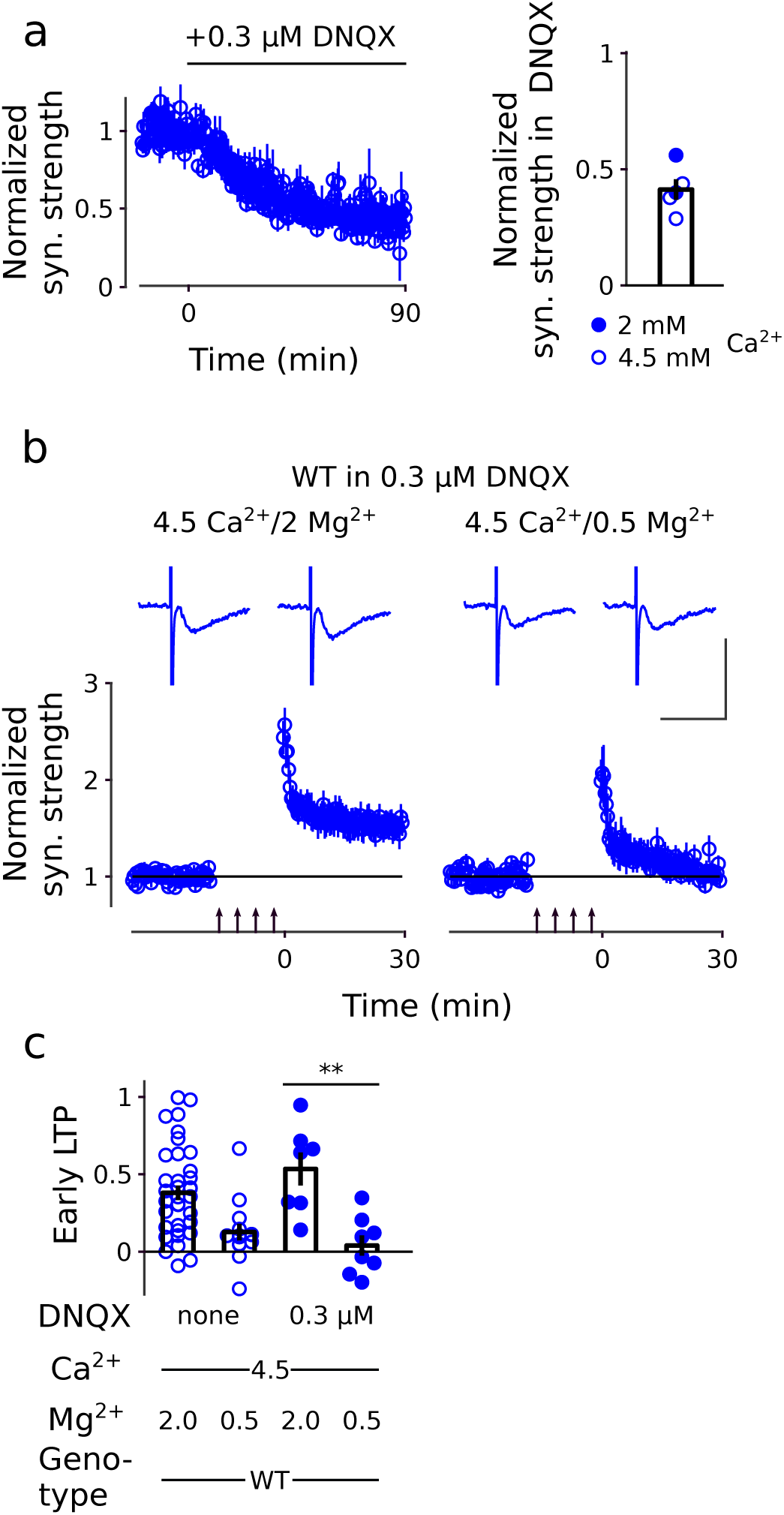
Early LTP remains occluded in 0.3 ➭M *DNQX*. **(a)** Response versus time when washing in 0.3 ➭M DNQX and relative response size between minutes 80 and 90 after DNQX application; mean was 0.41 *±* 0.05. Ca^2+^ was 4.5 mM for 3 of 5 experiments and 2 mM for the other 2. No substantial differences were seen and data were combined. Mg^2+^ was 2 mM throughout. Washing in DNQX was substantially slower than changing Ca^2+^ and Mg^2+^ levels (compare time course to time courses in Supplementary Figure S3). **(b)** Response versus time before and after tetanus (4 1 s-long trains at 100 Hz) in 0.3 ➭M DNQX (*n* ≥ 7). Ca^2+^ was 4.5, and Mg^2+^ was either 2 or 0.5 mM as indicated. Traces in insets are examples from before and 30 min after the 100 Hz trains; scale bars are 1 mV by 20 ms. **(c)** Quantification of potentiation 25-30 min after tetanus. The values for the first two bars are replicated from Fig 3 and are included for comparison. ** signifies *p <* 0.01, rank sum.

**Supplementary Figure S8.**
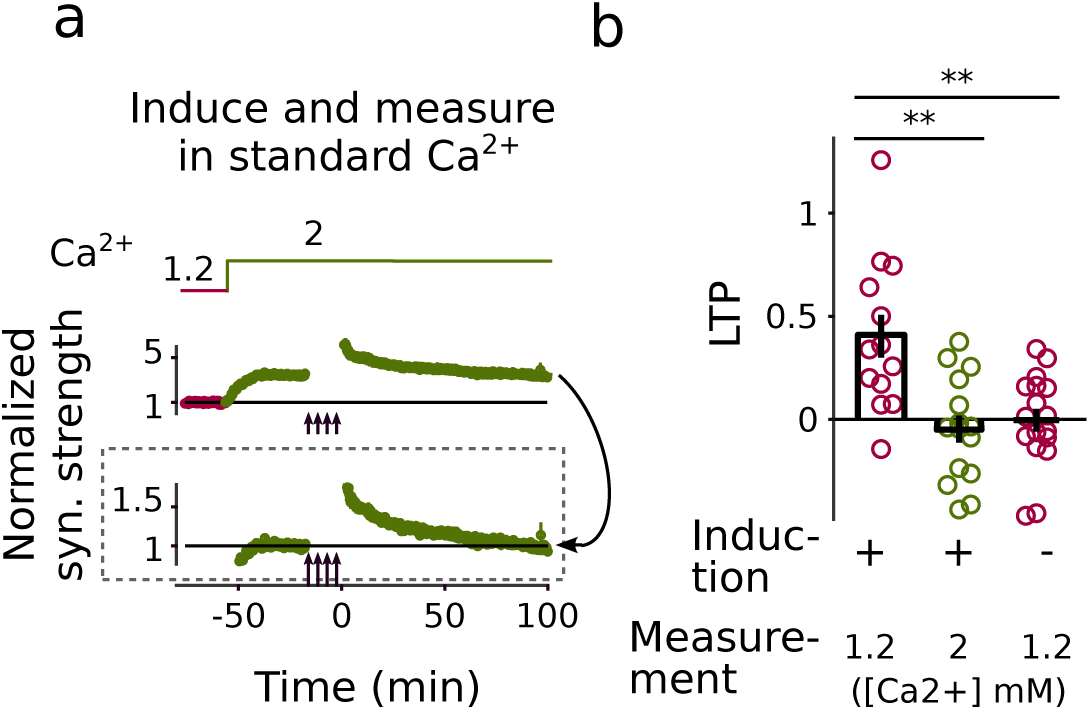
Interleaved control trials for Fig 4a. These trials confirm that the longest-lasting component of LTP is eliminated at SypFamilyKO synapses under standard conditions; i.e., when Ca^2+^ is maintained at 2 mM after induction. **(a)** Time course (*n* = 17). **(b)** The 2^2nd^ bar summarizes quantification of LTP when Ca^2+^ was maintained at 2 mM after LTP induction. Bars 1 and 3 are replicated from Fig 4a (** signifies *p <* 0.01, rank sum).

**Supplementary Figure S9.**
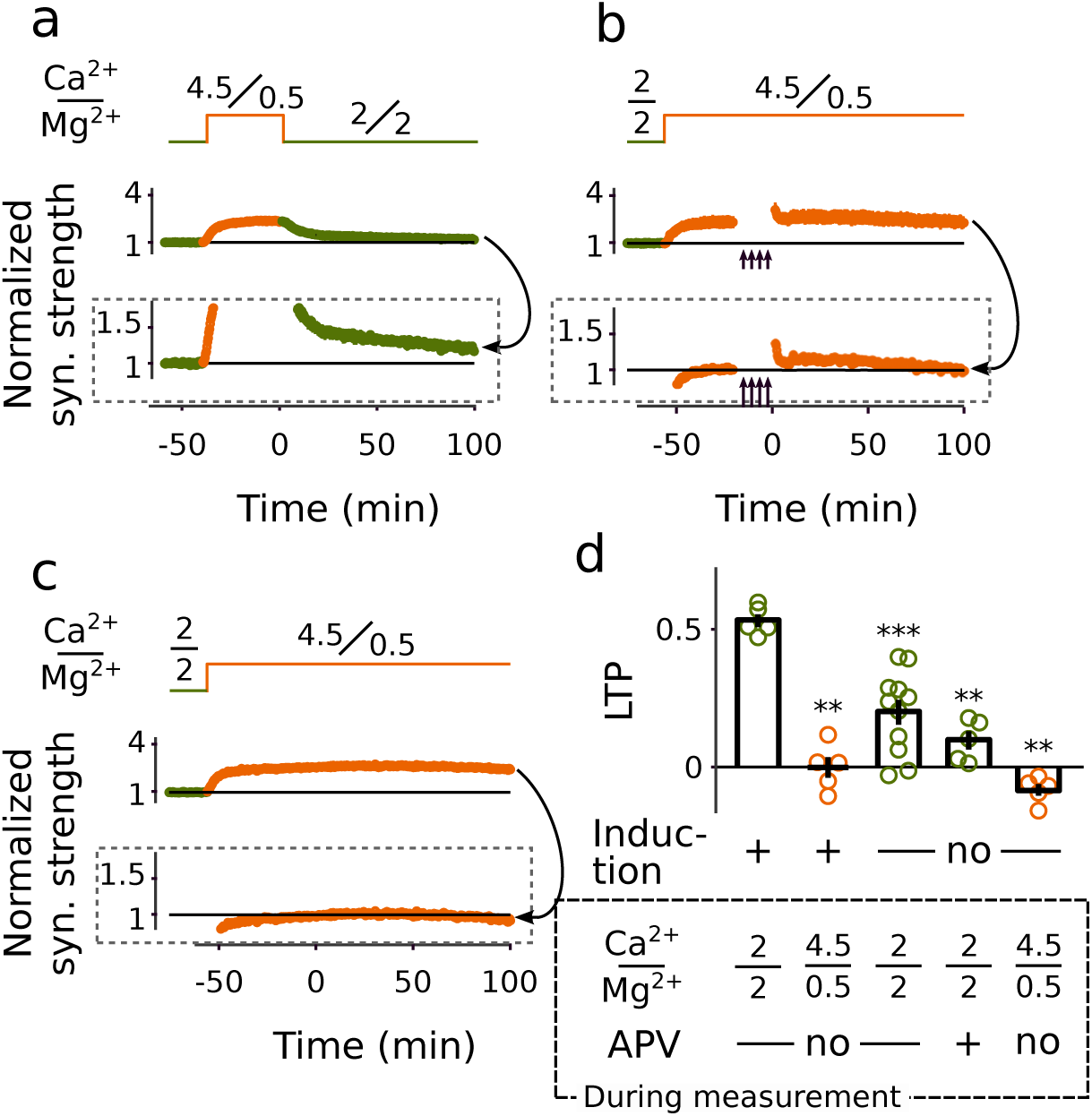
Interleaved control trials for Fig 4b. **(a)** Control trials where LTP was not induced before switching Ca^2+^/ Mg^2+^ back to 2/2 mM. These trials show that response size did not completely return to baseline in the absence of APV. Only 4 of 11 trials were interleaved with the trials from Fig 4b where LTP was induced, and the other 7 were conducted as follow-on experiments to confirm that responses did not fully return to baseline. **(b)** Controls where Ca^2+^/ Mg^2+^ was maintained at 4.5/0.5 mM after inducing LTP with 100 Hz stimulation, confirming that both the early and long-lasting components of LTP are eliminated when Ca^2+^/ Mg^2+^ is 4.5/0.5. All trials were interleaved with the trials in Fig 4b where LTP was induced. **(c)** Controls where Ca^2+^/ Mg^2+^ was maintained at 4.5/0.5 without inducing LTP with 100 Hz stimulation, confirming that responses are stable for hours when Ca^2+^/ Mg^2+^ is 4.5/0.5. All trials were interleaved with the trials in Fig 4b where LTP was induced. **(d)** Bars 1 and 4 are replicated from Fig 4b (** signifies *p <* 0.01, *** signifies *p <* 0.001, rank sum, *n ≥* 5).

## Notes

### Competing Interest Statement

The authors have declared no competing interest.

### Summary of Updates

The new version now shows that the early component of LTP can be occluded by raising the probability of release, whereas the previous version only showed this for the longest lasting version. In addition, we now include new experiment showing that LTP is blocked at the level of expression whereas maintenance and induction are both intact. The previous version only showed that induction was intact.

